# Exposure to maternal high-fat diet induces extensive changes in the brain of adult offspring

**DOI:** 10.1101/2020.07.17.209551

**Authors:** Darren J. Fernandes, Shoshana Spring, Anna R. Roy, Lily R. Qiu, Yohan Yee, Brian J. Nieman, Jason P. Lerch, Mark R. Palmert

## Abstract

Maternal environmental exposures, such as high-fat diets, diabetes and obesity, can induce long term effects in offspring. These effects include increased risk of neurodevelopmental disorders (NDDs) including autism spectrum disorder (ASD), attention deficit hyperactivity disorder (ADHD), depression and anxiety. The mechanisms underlying these late-life neurologic effects are unknown. In this article, we measured changes in the offspring brain and determined which brain regions are sensitive to maternal metabolic milieu and therefore may mediate NDD risk. We showed that mice exposed to maternal high-fat diet display extensive brain changes in adulthood despite being switched to low-fat diet at weaning. Brain regions impacted by early-life diet include the extended amygdalar system, which plays an important role in reward-seeking behaviour. Genes preferentially expressed in these regions have functions related to feeding behavior, while also being implicated in human NDDs, such as autism. Our data demonstrated that exposure to maternal high-fat diet in early-life leads to brain alterations that persist into adulthood, even after dietary modifications.

## Introduction

The prevalence of overweight/obesity is increasing among women of reproductive age[1][2]. Over-weight or obese mothers may predispose their children to adverse health outcomes – such as diabetes, obesity, and coronary heart disease[3][4][5]. More recently, the maternal environment has been recognized as an important factor in offspring brain development[6][7][8]. Evidence has shown that the perinatal exposure to maternal obesity and associated metabolic abnormalities increases the risk of neurodevelopmental disorders (NDDs) in offspring, including attention deficit hyperactivity disorder, autism spectrum disorders, anxiety, depression, schizophrenia, and eating disorders[9][10][11][12].

Due to the slow emergence of NDDs in humans, it is difficult to study mechanisms driving the association between maternal environment and NDDs in offspring. Mouse models provide an important opportunity to explore this link; the effects of typical human “Western” diets (which consists of 30-40% of calories from fats), including characteristic metabolic abnormalities, have been successfully recapitulated in mice fed diets consisting of 40-60% fat[13][14][15]. Furthermore, offspring of rodents fed high-fat diet exhibit a wide range of behavioural changes consistent with NDDs[16][17]. For example, rodents exposed to high-fat diet before weaning displayed increased susceptibility to depression-like behaviours[18], increased anxiety[19], and impaired spatial learning[20].

While the relationship between maternal diet and offspring NDD risk is likely to be mediated by developmental changes in particular brain structures, a systematic measure of alterations across the whole brain has not previously been reported. Since mice and humans have many homologous brain structures, the brain regions affected by maternal diet in mice may identify relevant structures in humans. Viral labelling of hippocampal neurons has already shown that dendritic arborization is impaired in offspring of mice fed high-fat diet[20]. However, the extent to which the rest of the brain may also be altered is unclear. Magnetic resonance (MR) imaging allows for the study of the whole brain in a high-throughput manner with sufficient resolution to quantify regional changes in volume or morphology[21].

Our aim was to explore the effects of maternal diet on offspring brain structure. Adult mice were fed one of three diets during breeding, gestation, and lactation: a low-fat diet with 10% of calories from fat (LF10) or high-fat diets with 45% or 60% of calories from fat (HF45 and HF60, respectively). After weaning, all offspring were raised on the low-fat diet until postnatal day 65 (P65), corresponding to early adulthood. MR imaging was used to measure neuroanatomical changes in adult offspring attributable to pre-weaning high-fat diet exposure. The Allen Brain Institute’s gene expression atlas was then used to identify which genes had spatial expression patterns consistent with the affected neuroanatomy.

## Results

### Dams fed high-fat diet showed significant changes in physiology but not brain structure

To verify effectiveness of the dietary manipulation, dams’ body weight, body composition, and glucose tolerance were measured. Figure 1 summarises how dams on high-fat diets (HF45 and HF60) showed the expected metabolic changes compared to dams fed the low-fat diet (LF10). Body weights of the different groups, though similar at the start of the study, began to diverge when the high-fat diet was introduced (Figure 1A). Dams fed a HF60 diet were significantly heavier than LF10 dams after approximately 2-3 weeks of acclimation. After 6 weeks, the dams acclimated to HF45 and HF60 diets had higher body fat percentage than LF10 dams (Figure 1B). Furthermore, dams on the HF60 diet showed abnormalities in the glucose tolerance test (Figure 1C).

**Figure 1:**
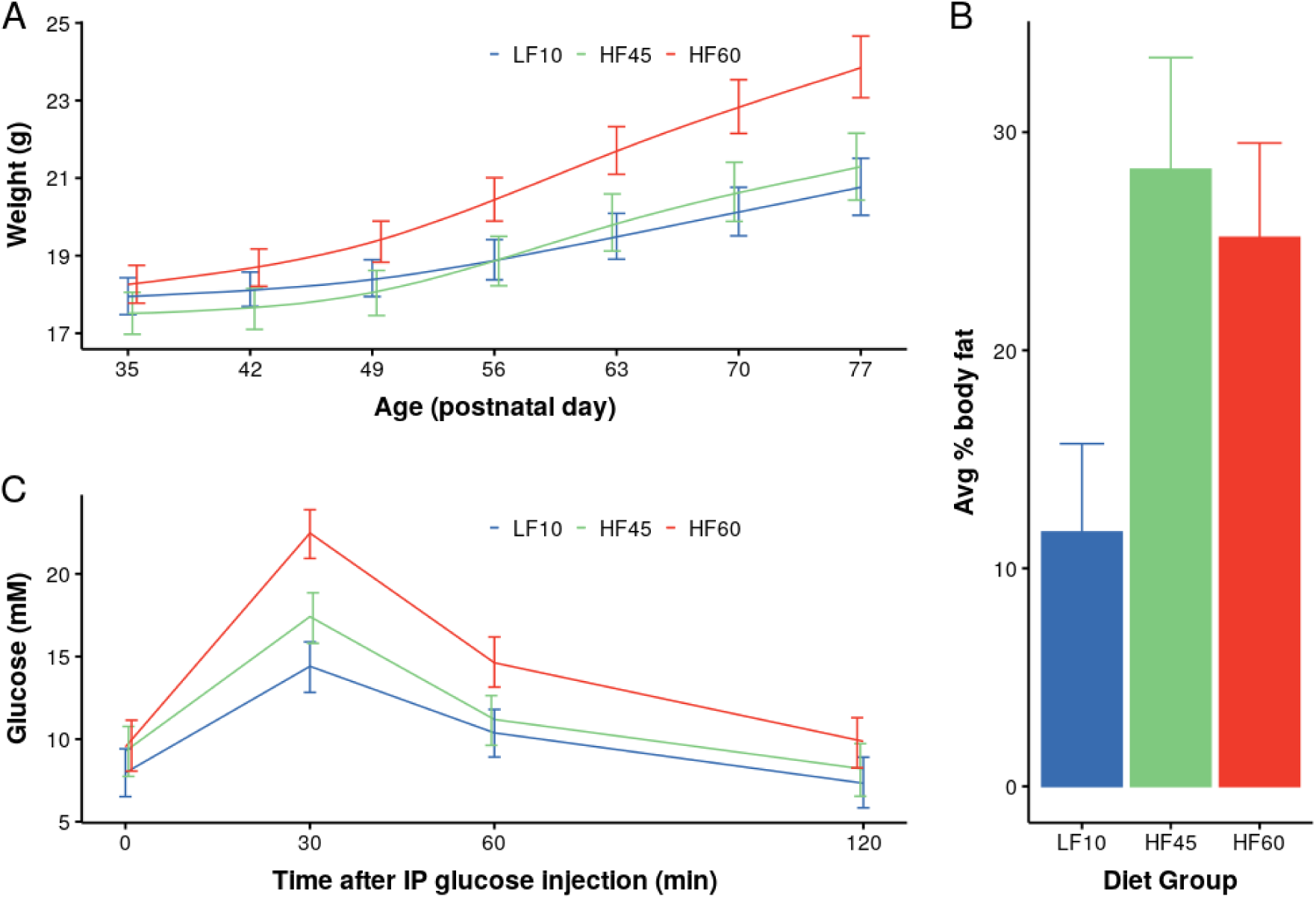
High-fat (HF) diet induced metabolic changes in dams. Mice were fed either a low-fat diet (LF10) or one of two high-fat diets (HF45 and HF60) during a 6-week acclimation period. (A) Body weights measured weekly were significantly affected by diet 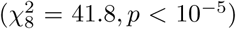, primarily driven by a diet-time interaction 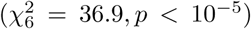. A significant difference between LF10 and HF60 groups emerged at around 2-3 weeks after the experimental diet was introduced. (B) Average body fat percentage after the 6-week acclimation period. There were significant differences in body-fat percentage between LF10 and HF45 diets (difference=16.6%, *p* < 10^−3^), and LF10 and HF60 diets (difference=13.5%, *p* < 10^−3^); and no significant differences between the HF45 and HF60 diets (difference=3.1, *p* = 0.6). (C) Intraperitoneal (IP) glucose tolerance tests were performed after the 6-week diet acclimation period. There was a significant difference in glucose levels 30 minutes after IP injections (difference = 8.04mM, *p* < 10^−8^) and 60 minutes after IP injections (difference = 4.25mM, *p* < 10^−2^) between the LF10 and the HF60 groups. All bars represent 95% confidence intervals.

After the 6-week acclimation period, all dams (LF10, HF45, HF60) continued on their respective diets throughout breeding, gestation, and lactation until the end of weaning – resulting in approximately 12 weeks of dietary exposure. Subsequently, dams were sacrificed in order to image their brains. While high-fat diet had an impact on physiology, it did not significantly affect neuroanatomy of dams (Figure 2). Voxel-wise statistics, illustrated in Figure 2A, show this to be the case throughout the brain. Of the 182 bilateral structures in our atlas, none of the structure volumes were significantly affected by dietary fat percentage (examples shown in Figure 2C,D). The total brain volume was also not impacted significantly by diet (Figure 2B).

**Figure 2:**
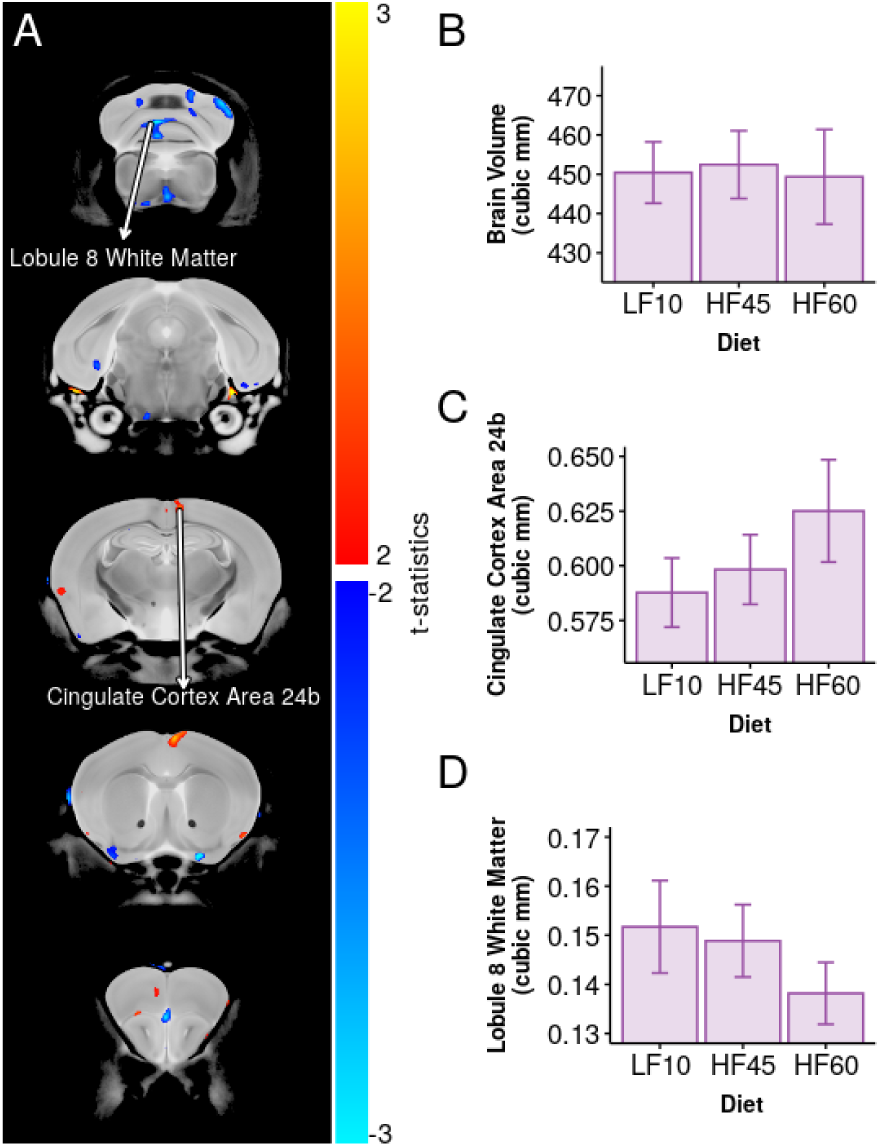
High-fat diet did not have a significant effect on the brain structure of dams. Trends in associations of dietary fat-percentage with decreases (blue/turquoise colours) and increases (red/yellow colours) in the measured volume of brain regions were not significant. (B) The whole brain volume was not significantly affected by diet (*F*_1,22_ = 0.003, *p* = 0.96). One example structure is the cingulate cortex area 24b (C) which was positively correlated with dietary-fat percentage (*t*_22_ = 2.5, *p* = 0.02, FDR = 0.92, not significant). Another example structure is the white matter of cerebellar lobule 8 (D) which showed decreased volume with increased fat-percentage (*t*_22_ = −2.13, *p* = 0.045, FDR = 0.98, not significant).

### Metabolic effects of maternal diet were ameliorated after high-fat diet was substituted with low-fat diet

To limit experimental diet exposure to the *in utero* and perinatal period, offspring from all diet groups were placed on low-fat diet at weaning on postnatal day 21 (P21). By early adulthood (P65), the change to the low-fat diet resulted in attenuation of all initial per group differences revealed in our physiological measurements (Figure 3). Figure 3A shows that offspring fed high-fat diets in early-life had significantly greater body weight than those fed low-fat diet at P21. However, after two weeks of being weaned onto a low-fat diet, all diet groups had overlapping 95% confidence intervals. Similarly, offspring exposed to high-fat diet in early-life did not have significantly different body fat composition at P65 compared to offspring raised solely on low-fat diet (Figure 3B). Upon intraperitoneal administration of glucose, there was no abnormal elevation of blood glucose concentration in HF45 and HF60 adult offspring (Figure 3C), unlike their mothers (Figure 1C). There was a decrease in blood glucose concentration 30 minutes after IP glucose administration in male HF45 offspring compared to male LF10 offspring (SR_96.8,24_ = 4.36, *p*_*tukey*_ < 0.01, SR is the Studentized Range distribution). No such decrease was found between the HF60 and LF10 male offspring.

**Figure 3:**
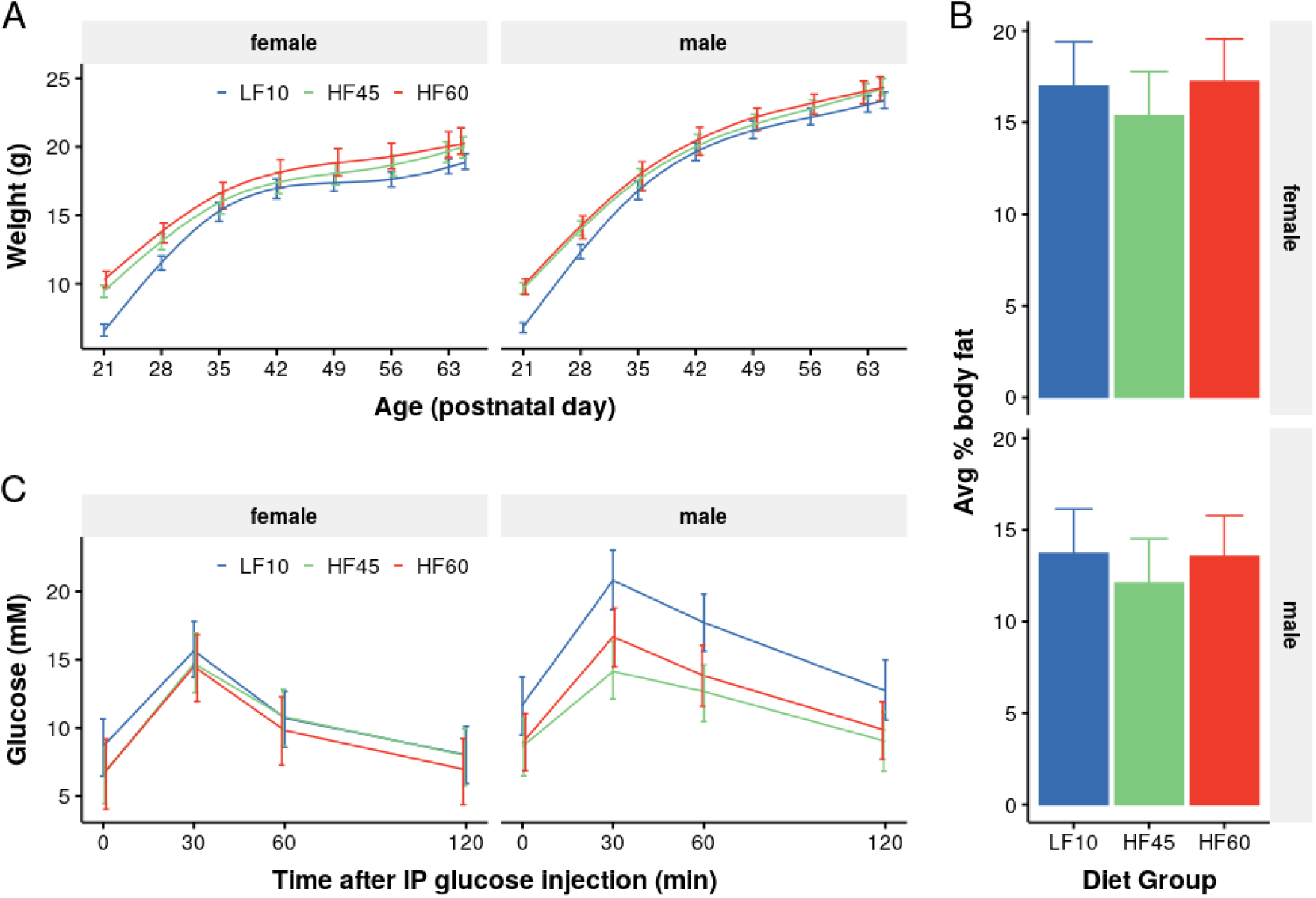
Offspring exposed to maternal high-fat diets showed amelioration of metabolic effects when weaned to a low-fat diet. Prior to weaning (P21), offspring were exposed to either the low-fat diet (LF10) or one of two high-fat diets (HF45 and HF60). (A) Body weights of HF45 and HF60 mice normalized after being weaned onto the low-fat diet. Body weight trajectories had a significant diet-time interaction 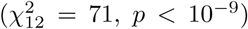. At weaning (P21), the LF10 female offspring were 6.6g [6.2,7.0] and male offspring were 6.8g [6.4,7.3], where square brackets represent 95% confidence intervals. HF45 offspring weighed significantly more with females weighing 9.5g [9.0, 10.0] and males weighing 9.6g [9.2,10.0]; HF60 offspring were also significantly heavier (females 10.3g [9.7,10.9], males 9.9g [9.3,10.5]). After 2 weeks, with all mice weaned onto the same low-fat diet, the body weights of the different groups were similar (LF10 females 15.3g [14.6,15.9], HF45 females 15.8g [15.1,16.7], HF60 females 16.5g [15.5,17.5], LF10 males 16.8g [16.1,17.5], HF45 males 17.5g [16.9,18.2], HF60 males 17.9g [17.0,18.9]). (B) Body-fat composition was measured in early adulthood (P65) and there were no significant differences between the different diet groups (*F*_2,63_ = 1.20, *p* = 0.3). (C) Glucose tolerance results in adulthood were also not significantly different among the diet groups at P65. There was neither a significant effect of early-life diet 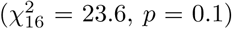 nor a significant diet-time interaction 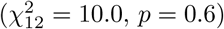. All bars represent 95% confidence intervals.

### High-fat diet in early-life programmed brain structure changes in adulthood

Offspring were sacrificed in early adulthood (P65) to image neuroanatomy. Their brains showed widespread volume changes in several regions due to high-fat diet exposure during gestation and lactation (Figures 4 and 5). These changes were present despite offspring transition to low-fat diet for 6 weeks post-weaning and amelioration of physiological abnormalities (Figure 3). The widespread effect of high-fat diet exposure on brain structure can be seen in Figure 4A. The volume of the whole brain was impacted by the early-life diet (Figure 4C). The effect of diet was found to be significantly different among males and females in a small number of isolated areas, such as voxels in the medial amygdala and the basal forebrain (Figure 4B). However, these sex-interactions were only significant in subregions of these structures and were not considered significant when analyzing the bilateral structures as a whole (Figure 4D,E). Volume and effect-sizes for all brain structures are reported in Appendix Table 1.

**Figure 4:**
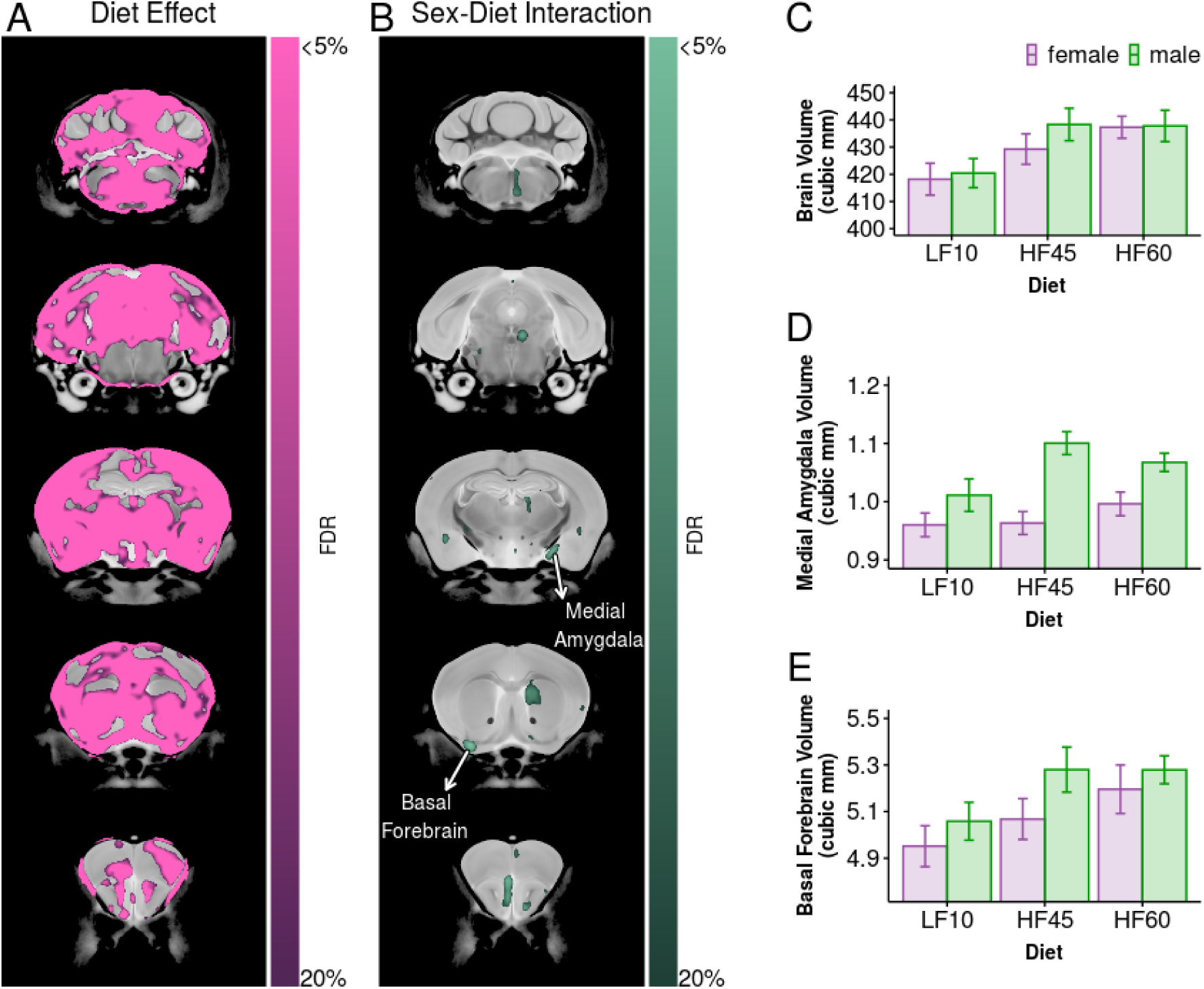
Offspring exposed to early-life high-fat diet exhibit extensive structural brain changes in adulthood. (A) After conducting an ANOVA at every voxel, early-life high-fat diet was found to affect the volume of several regions throughout the brain. (B) In some brain regions, the effect of diet was modulated by sex. (C) Much of the effect of diet was related to overall brain size (*F*_2,95_ = 24, *p* < 10^−8^). Two important structures in the extended-amygdalar network are the medial amygdala and the basal forebrain. (D) Absolute volumes of the medial amygdala and the basal forebrain were strongly impacted by diet (*F*_2,95_ = 13, *p* < 10^−4^, FDR < 10^−4^ and *F*_2,95_ = 14, *p* < 10^−5^, FDR < 10^−4^, respectively). While some parts of right medial amygdala had a significant sex-diet interactions (shown in B), the medial amygdala as a whole did not have significant sex interactions (*F*_2,95_ = 8.9, *p* < 10^−3^, FDR = 0.051). A significance threshold of 5% FDR is shown by the saturated colours (pink for the effect of diet and teal for the diet-sex interaction). The colour bar extends to 20% FDR to allow visualization of cluster extent. Error bars represent 95% confidence intervals. Structure volume and effect-sizes are reported in Appendix Table 1.

**Figure 5:**
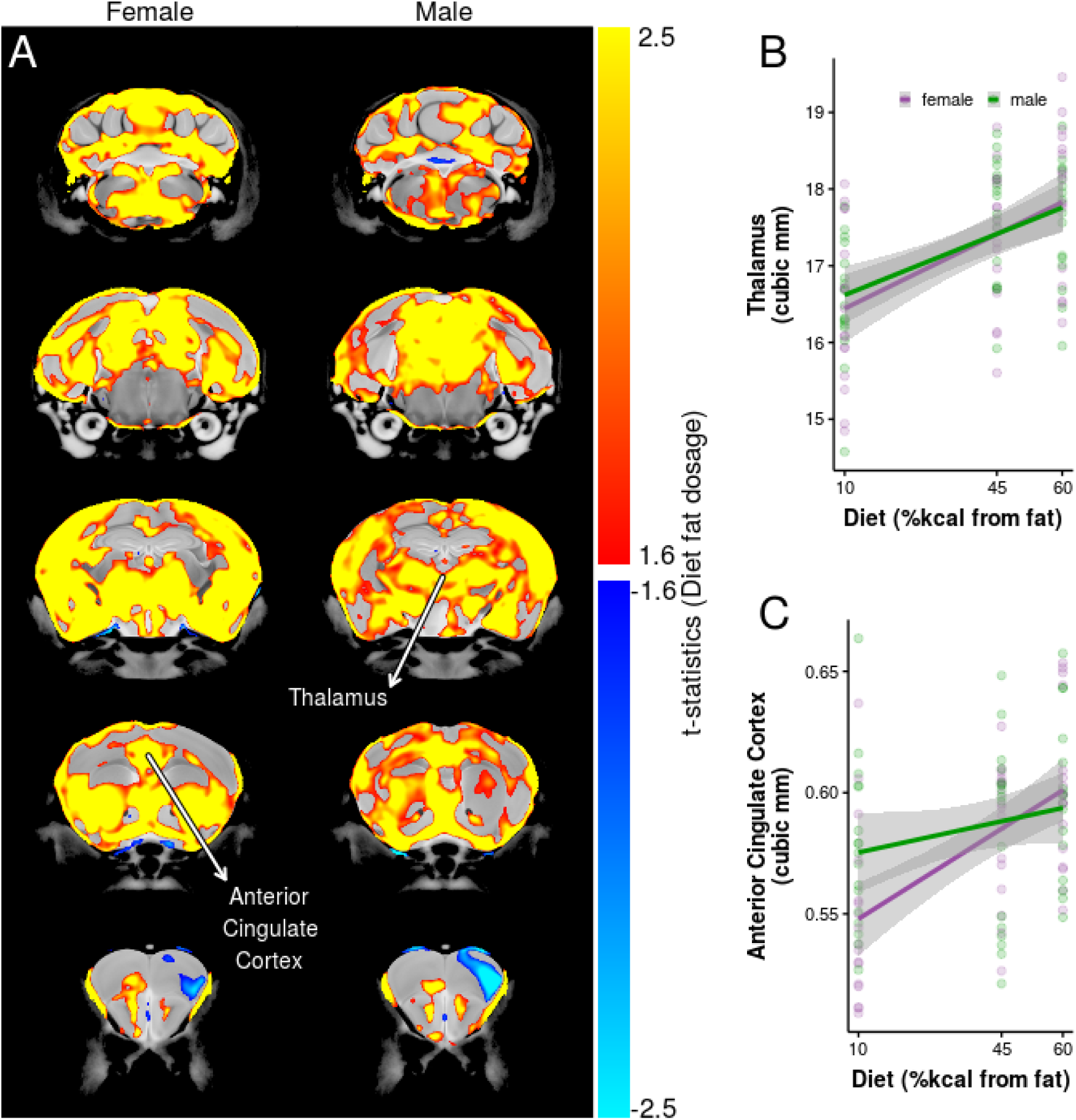
Fat content of early-life diet correlated with increased volume of several brain regions. (A) Brain regions showed widespread increases in measured volume as dietary fat percentage increased for both males and females (red/yellow colour scale). Few regions, such as the primary motor cortex, showed a volume decrease (blue/turquoise colour scale). The volumes of the (A) thalamus and (B) anterior cingulate cortex correlated with early-life dietary fat percentage for both males and females. A significance threshold of 5% FDR is shown by the extremes in the colour bar for positive (yellow) and negative (turquoise) dosage effects; the colour bar extends to 20% FDR (red/blue) to allow visualization of cluster extent. Shaded regions in (B) represent 95% confidence intervals. Structure volume and effect-sizes are reported in Appendix Table 1.

Since our study included 2 high-fat diets with differing fat percentage, we were able to explore possible dose-response effects. We interrogated the effect of diet on the brain by examining whether the volume of brain regions increased or decreased with dietary fat-percentage (Figure 5). Consistent with Figure 4, dietary fat-percentage had a widespread positive correlation with the volume of several brain regions (Figure 5A). The thalamus and anterior cingulate cortex are shown as examples (Figure 5B,C).

### Brain regions sensitive to early-life high-fat diet were enriched with genes involved with behaviour and NDDs

Maternal diet had a significant impact on the anatomy of several brain regions in the adult mouse. To find candidate genes associated with the observed brain changes, we registered our MR data to the Allen Brain Institute (ABI) gene expression atlas[22]. Shown in Figure 6B (first image row), we defined a Region-Of-Interest (ROI) as brain areas where early-life high-fat diet had a highly significant impact on neuroanatomy (FDR < 0.01). We then used spatial gene expression analysis[23] to identify genes that had a spatial expression pattern consistent with this ROI. Candidate genes were identified by preferential expression in the ROI (i.e. higher expression within the ROI and reduced expression outside the ROI). Preferential expression was quantified by a fold-change measure; average gene expression in ROI divided by average gene expression in the whole brain. This fold-change measure is greater than 1 when a gene has higher expression in diet-sensitive brain regions than its brain-wide average. The ABI adult gene expression atlas spans the whole mouse genome, but a small subset of genes had their expression patterns quantified throughout development. For this subset of genes, we also considered how gene expression patterns in our ROI changed during postnatal development. We found 3 main clusters of genes: one were expression in ROI was low during neonatal life and increased during the juvenile period (P28), one were expression was high during neonatal life and decreased during the juvenile period (P28), and one where expression was high during neonatal life and decreased before weaning (P14). The adult and development gene expression data are summarised in Appendix Table 2.

**Figure 6:**
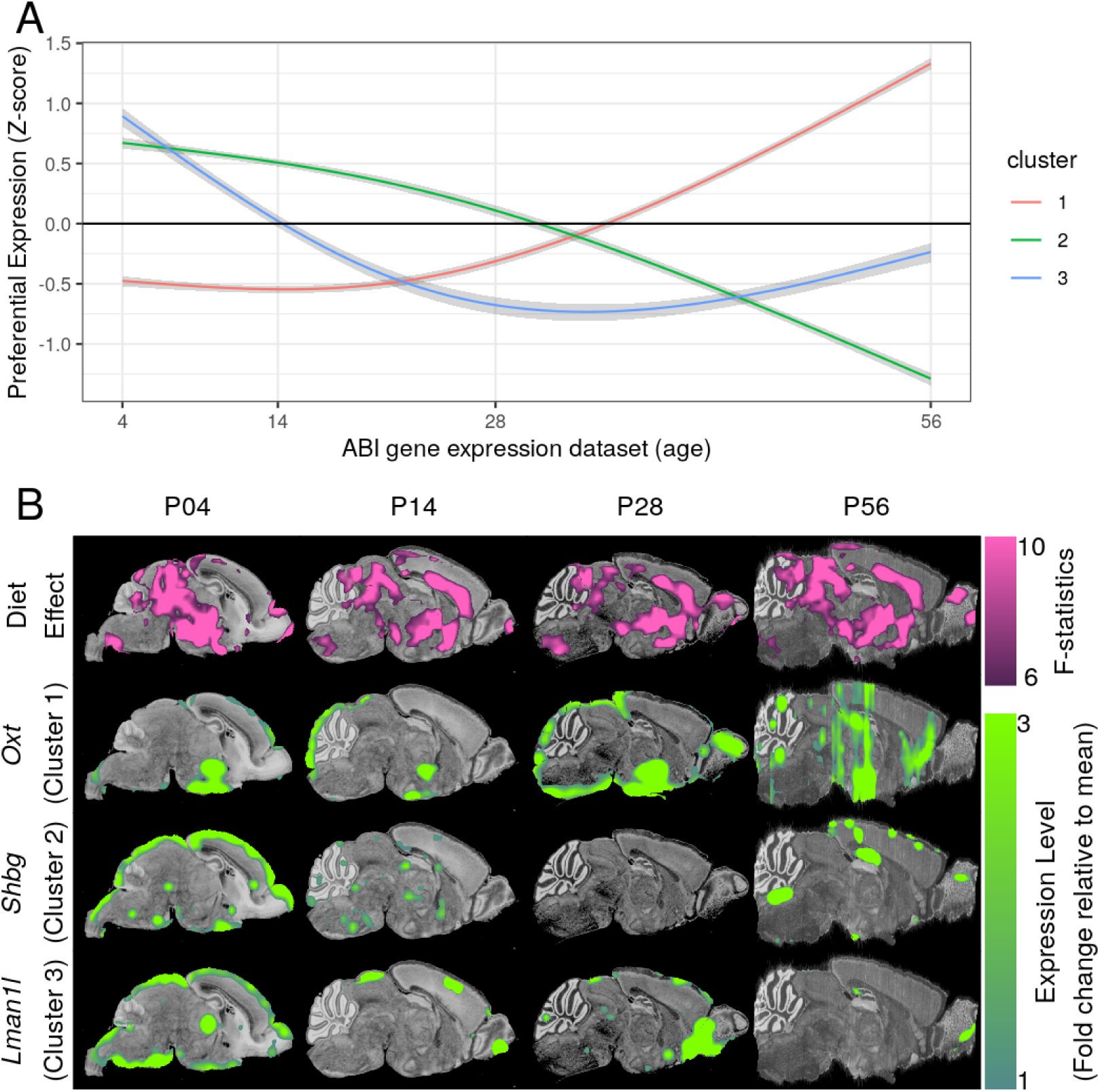
Genes expressed in brain-regions sensitive to early-life diet over the course of mouse development. Brain regions sensitive to the early-life diet constituted the ROI for spatial gene expression analysis and were defined as regions where early-life diet had a highly-significant effect (FDR < 0.01) on adult neuroanatomy. The ROI was transformed to the nissl atlases defining the ABI gene expression dataset (first image row of B) and the spatial expression of genes were quantified throughout the course of postnatal development. Shown in A, three clusters of spatio-temporal gene expression were found: cluster 1 showed genes that had low expression in ROI during neonatal life and increased during juvenile life, cluster 2 showed genes that had high expression in ROI during neonatal life and decreased when subjects become juveniles, and cluster 3 showed genes that had high expression in ROI during neonatal life and decreased rapidly before weaning. *Oxt* was chosen as a representative example of a gene associated with ASD and part of Cluster 1 (image row 2 in B). *Shbg* had the highest expression in Cluster 2 (image row 3 in B) and *Lman1l* had the highest expression in Cluster 3 (image row 4 in B). Shaded-regions represent 95% confidence intervals, estimated by fitting a linear mixed-effects model to the data (fixed effects of quadratic natural spline for age, cluster membership, and all interactions, random intercept for every gene).

Spatial gene expression analysis revealed candidate genes, whose spatial expression pattern was similar to the pattern of change induced in offspring brains by maternal high-fat diet. The development gene expression data only consists of 2107 genes, making it unreliable for popular automated analysis techniques like Gene Ontology (GO) enrichment analysis. Instead, we considered the genes with the highest expression in each cluster individually. Sex-hormone-binding-globulin (*Shbg*) had the highest expression in Cluster 2 and Lectin-Mannose-Binding-1-Like (*Lman1l*) had the highest expression in Cluster 3 (Figure 6B third and fourth image row).

Genes in Cluster 1 have high expression in diet-sensitive brain regions during adulthood. Unlike the ABI developmental dataset, the ABI adult gene expression data spanned the whole mouse genome. Thus, it was possible to conduct GO enrichment analysis to identify functions and disease processes associated with the preferentially-expressed genes. We ran a GO enrichment analysis on the top 4000 genes with the highest preferential expression in the adult brain using the `GORilla` tool[24], the results of which are included as Appendix Table 3. Significant GO terms included *synaptic signalling* (GO:0099536, *q* = 0.03), *feeding behavior* (GO:0007631, *q* = 0.02), and *neuron differentiation* (GO:0030182, *q* < 0.01). Similar results were also seen with a ranked-list of all genes using the minimum-hypergeometric test.

Using the Mouse Genome Informatics (MGI) database, we examined disease processes associated with genes that had spatial expression similar to our identified ROI. The MGI database has human diseases annotated with human genes and their homologous mouse genes. For every annotated human disease, we found the homologous mouse genes and determined their average fold-change. A high average fold-change implies that these genes had greater expression in the ROI (defined as diet-sensitive brain regions) relative to the rest of the brain. Bootstrap sampling was used to assess significance and Bonferroni correction was used to account for multiple-comparisons across the diseases. Genes associated with ASD had significantly higher preferential expression than the background set of all genes (*p*_*adjusted*_ = 0.017) and was the only significant disease term. The next most significant disease terms were schizophrenia and hydrocephalus (*p*_*uncorrected*_ = 0.01 and 0.02, respectively) but they were not considered significant after correction for multiple comparisons (*p*_*adjusted*_ = 0.15 and 0.28, respectively).

We sought to validate our findings that brain regions sensitive to early-life diet display preferential expression of mouse genes whose human homologues are associated with ASD. The Simons Foundation Autism Research Initiative (SFARI) database was queried to obtain a list of single-gene-mutant mouse models of autism. We found the average fold-change in these genes, and compared it to the average fold-change in all genes, estimating significance using bootstrapping. Similar to the analysis with the MGI dataset, genes associated with mouse models of autism had significantly higher preferential expression in brain regions sensitive to early-life diet (*p* < 10^−3^). For visualization, *Oxt* was chosen as a representative of ASD-associated gene with increased spatial preferential expression (Figure 6B second image row).

## Discussion

We demonstrated that high-fat exposure in early-life (based on maternal diet during gestation and lactation) caused structural changes in offspring brains that persisted into adulthood, even after the high-fat diet was removed and metabolic abnormalities attenuated. These data, coupled with the lack of diet effect on maternal brain structure, suggests that brain development may be particularly vulnerable to effects of high-fat diet during early-life. While there have been studies showing the effect of high-fat diet on older, juvenile mice, these studies generally focus on particular brain structures and behaviours[25][26]. To our knowledge, our study is the first to examine the whole brain and show that many structures in the developing brain are vulnerable to effects of maternal/early-life diet.

As proof of principle, we replicated existing findings that mice fed a diet consisting of 40-60% fat develop physical and metabolic abnormalities[13][14][15], verifying that our diets and acclimation period had the desired effects on dams. This allowed us to model the metabolic effects in humans consuming high-fat diets. In our study, the high-fat diet caused a significant increase in body weight and body fat percentage in dams, as well as affected glucose tolerance. It is worth noting that while this type of diet is used extensively in literature to explore the effects of high-fat intake, the diet is correspondingly low in carbohydrate content. Thus, it is not possible to separate the effect of low carbohydrate intake from high-fat intake in this study. However, this aspect is likely not a substantial confound for our model given that increasing fat intake in humans tends to also decrease carbohydrate intake[27]. Furthermore, Huang et al. showed that varying carbohydrate and protein intake in mice fed high-fat diets did not significantly impact level of obesity[28].

It is also worth noting that we compared effects of high-fat diet to a low-fat diet (10% kcal from fat). While this control is used extensively in literature, it is different from the standard mouse facility diet containing 18% kcal from fat. This may explain the overall low-brain volumes seen in low-fat diet-fed offspring mice (Figure 4C) compared to their mothers (Figure 2D) and literature[29]. It may also explain differences between offspring and dams in body-fat composition (Figure 1B and Figure 3B).

We also found maternal brain anatomy was not substantially impacted by the high-fat diet given over the course of approximately 12 weeks. Our results, conducted on C57Bl6/J mice, were consistent with results from Rollins et al, who found that 16 weeks of high-fat diet (60% calories from fat) were needed before adult mice (C57Bl6/J x 129S1/SvImJ strain) had a neuroanatomical phenotype detectable by MR imaging[30].

Offspring exposed to maternal high-fat diet during the *in utero* and perinatal periods showed some recovery in metabolic measures after transitioning to the low-fat diet at weaning. This result is similar to other studies investigating how transition to a low-fat diet can help obese mice recover physiologic and metabolic functions[31]. The substantial metabolic recovery allowed us to minimize confounds of aberrant metabolism during adulthood that could affect brain structure. Thus, the brain structure changes seen in the adult mice were less likely due to ongoing metabolic abnormalities and, instead, more likely secondary to the longstanding effects of diet/metabolic milieu during *in utero* and perinatal life, which are important periods of organism development.

The advantage of using MR imaging in our study was that we could show the significant and widespread impact that early-life high-fat diet has on brain anatomy in later life. These brain alteration may provide insights towards the complex neural mechanisms responsible for the increased NDD risk in offspring exposed to maternal obesity. Additionally, we could also identify particular structures with known behavioural roles to generate hypotheses regarding how the impacted anatomy could influence organism behaviour.

The basal forebrain and medial amygdala are part of the extended-amygdala network and were found to be affected by early-life high-fat diet exposure. These structures have a well-known role in reward-seeking behaviours and have been investigated extensively in the context of drug addiction[32]. More recently, studies have shown that the basal forebrain may also have an important role in food-seeking behaviour[33]. Although not the focus of our studies, the later life effects we observed in these structures raises the possibility that analogous alterations may contribute to the increased rates of adverse health outcomes seen in human children born to overweight/obese mothers[3][4][5]. Pertinent to our study of NDD risk is the role these structures are thought to play in etiology of disorders such as ASD and anxiety[34]. The medial amygdala is also an important mediator of sex-specific behaviours[35] and is known to be sexually dimorphic[36]. In some parts of the medial amygdala, we observed that the effect of diet on region volume was significantly modulated by sex. However, this sex interaction failed to reach significance for the medial amygdala as a whole. This preliminary finding should be investigated further given the sex differences in rates of NDDs.

In our final analyses, we used the ABI gene expression atlas[22] to identify genes that are highly expressed in brain regions sensitive to early-life high-fat diet exposure. These candidate genes may drive the relationship between maternal diet and NDDs in offspring. The developing gene expression atlas revealed several genes that had high expression in these brain regions in neonatal life. Among these genes are *Shbg*, which plays an important role in modulating the effects of androgens and estrogens[37], and *Lman1l*, which is associated with schizophrenia[38].

The adult gene expression atlas spans the whole genome, making it reliable enough for GO enrichment analysis. We found that genes with the highest spatial preference in diet-sensitive brain regions in adulthood tend to be significantly involved with biological processes that impact organism feeding behaviour. Furthermore, genes associated with autism behaviours – in both mice and humans – were also found to have significantly higher spatial expression in brain regions impacted by early-life high-fat diet. These genes and related pathways may be important starting points for exploring genetic mechanisms linking maternal metabolic milieu to NDD risk in offspring.

## Conclusion

Exposure to high-fat diet *in utero* and during perinatal development had a significant impact on brain structure in adulthood. These effects were persistent despite a change to a low-fat diet that attenuated the metabolic effects of early-life high-fat diet exposure. The impacted brain regions included regions that play an important role in the neural circuitry of reward and memory and exhibited a high expression of genes associated with behaviour and ASD. Our results demonstrate that the link between early-life high-fat diet exposure and risk for NDDs is also associated with brain structure changes, and provide strong evidence that the mouse can be used to investigate this relationship further.

## Methods

### Mice

All animal experiments were approved by The Centre for Phenogenomics (TCP) Animal Care Committee (AUP 17-0175H) in accordance with recommendations of the Canadian Council on Animal Care (CCAC), the requirements under the Animals for Research Act, RSO 1980, and the TCP Committee Policies and Guidelines. All mice were bred and maintained under controlled conditions (25 °C, 10/14 h light/dark cycle) at TCP in sterile, individually ventilated cages and were provided irradiated chow and sterile water *ad libitum*.

### Maternal and offspring diet

Five-week old female and male C57Bl6/J mice raised on the Teklad global 18% protein diet (18% kcal from fat, 24% kcal from protein, 58% kcal from carbohydrate, diet 2918, Envigo, Indianapolis, IN) were acclimated to one of three new diets at the start of the study – a low fat diet with 10% kcal from fat (LF10), high-fat diet with 45% kcal from fat (HF45), or high-fat diet with 60% kcal from fat (HF60) for six weeks (see Supplementary Table 1 and 2 for detailed diet information). After diet acclimation, diet-fed-matched females and males were paired for mating. Impregnated females were individually housed and remained on their respective diets throughout breeding, gestation and lactation. The resulting offspring pups were weaned at P21 and group-housed with 4-5 animals per cage. All offspring mice were fed the LF10 diet after weaning and throughout adulthood.

### Growth rates, body composition and metabolic testing

Weekly weights of females were recorded throughout diet acclimation (18 mice for the LF10 diet, 15 mice for the HF45 diet, 18 mice for the HF60 diet). Offspring mice were weaned and also weighed regularly to assess growth patterns until P65 (37 females and 36 males from 8 dams fed LF10 diet, 24 females and 37 males from 8 dams fed HF45 diet, 15 females and 17 males from 6 dams fed HF60 diet).

A bench-top NMR analyzer (Minispec, Bruker, Bellerica, MA) was used to analyze whole body composition and to distinguish fat tissue from lean tissue and free fluids *in-vivo* as we have done previously [39]. Maternal mice were assessed for body composition after diet acclimation but prior to breeding, when they were approximately 11 weeks of age (18 mice for LF10 diet, 15 mice for HF45 diet, 18 mice for HF60 diet). Offspring mice were assessed for body composition at approximately 9 weeks of age (11 females and males for the LF10 and HF45 diets, 12 females and 13 males for the HF60 diet).

Glucose tolerance testing (GTT) was conducted based on previously published protocols[39][40]. Briefly, mice were housed without food for 6 hours. Blood glucose levels were checked and then a 10% glucose solution was administered via intraperitoneal (IP) injection at a dose of 0.01 mL/g. Blood glucose levels were checked again 30, 60 and 120 minutes post injection. GTT was conducted on dams at the end of the diet acclimation period at 11 weeks of age (12 mice for each diet group) as well as in adult offspring at 9 weeks of age (8 females and males for the LF10 and HF45 diets, 6 females and 8 males for the HF60 diet).

### Brain perfusion, fixation, and image acquisition

Fixed brain samples were prepared for MR imaging to assess neuroanatomical changes of dams (8 mice per diet group) and offspring due to maternal/early-life diet (17 females and males for the LF10 and HF45 diets, 17 females and 16 males for the HF60 diet). Two of the 8 dams in the HF60 group did not give birth (either reabsorbed the pregnancy or cannibalized the litter without evidence of birth), but were still sacrificed and imaged. A previously described fixation protocol[41] was conducted on P127 ± 9 dams and P65 ± 3 offspring (median ± max range). Briefly, mice were anesthetized and placed in a supine position with thoracic cavities opened allowing for access to the heart. Perfusion occurred through the left ventricle with 30 mL of phosphate-buffered solution (PBS) (pH 7.4), 2 mM Prohance^®^ (gadoteridol, Bracco Diagnostics Inc., Princeton, NJ) and 1 *µ*L/mL heparin (1,000 USP units/ml, Fresenius Kabi Canada Ltd., Toronto, ON) at room temperature (25 °C) at a rate of approximately 60 mL/h. This was followed by fixation with 30 mL of 4% paraformaldehyde (PFA) in PBS containing 2 mM ProHance^®^ at the same rate. Following fixation, the heads were removed from the bodies along with the skin, ears, and lower jaw. The skull structure was allowed to postfix in the fixation solution at 4 °C for approximately 12 h. Samples were then placed in a solution of PBS, 2 mM ProHance^®^, and 0.02% sodium azide (sodium trinitride, Fisher Scientific, Nepean, ON) at 4 °C for storage until imaging. Brains were imaged at least 2 weeks postmortem.

A multi-channel 7.0-T scanner with a 40-cm diameter bore magnet (Varian Inc., Palo Alto, CA) was used to acquire three-dimensional (3D) anatomical images. A custom-built 16-coil solenoid array was used to image 16 brain samples concurrently[21]. In preparation for imaging, the samples were removed from the storage solution, blotted and placed in 13mm-diameter plastic tubes filled with a proton-free susceptibility-matching fluid (Fluorinert FC-77, 3M Corp. St. Paul, MN). A neuroanatomical scan was performed using a T2-weighted, 3D fast spin echo sequence using a cylindrical k-space acquisition with TR=350ms, TE=12ms, ETL=6, two averages, FOV (mm) 20×20×25, and matrix size of 504×504×630[42]. Total imaging time was 14 hours and resulted in forty-micron isotropic resolution images of brains within skulls. To correct for geometric distortions, MR images of precision-machined phantoms were aligned towards a computed tomography (CT) scan of the same phantom to produce transformations in a coil-specific manner.

### Image registration

To quantify the anatomical differences between images, all of the 24 maternal brains and 101 offspring brains were registered together. This procedure brought images into a precise yet unbiased anatomical alignment in a multi-step process and has been previously described[43]. First, the `mni autoreg` tools[44] were used to globally align images using a series of affine transformations (rotations, translations, scales, and shears). Then, the Advanced Normalization Tools (ANTS)[45] were used to locally deform images through an iterative process to achieve optimal alignment.

The pydpiper toolkit[46] was used to create a pipeline that performed the image registrations in an automated fashion. The output of the pipeline was a population average which represents the average anatomy of all the mice included in the study and transformations that mapped the average to the individual images. The MAGeT[47] pipeline was then used to automatically segment the 125 brain images into 336 structures using a previously published segmentation[48][49][50][51][52]. Summing the volume of bilateral structures resulted in 182 total structures. At the end of the registration pipeline, every MR image had an associated transformation which governed how the population average had to deform to align to the MR image. The jacobian determinant of this transformation quantifies the volumetric change at each voxel. To eliminate skewness inherent to the jacobian determinants, the log of the result was computed, which was then statistically analyzed to quantify region-specific neuroanatomy phenotype.

### Data analysis

All statistical analyses were conducted in the `R` software environment. Data manipulations and visualizations were done using the `tidyverse` suite of packages[53]. All linear mixed-effects models were fitted using maximum-likelihood and model-fitting was performed using the `lmerTest` package[54][55].

### Growth rates, body composition and metabolic testing

Maternal and offspring body weight data were analysed with linear mixed-effects models. For maternal body weight measurements, the model contained fixed effects of diet as a categorical variable, of time-on-diet as a cubic natural spline, and all interactions. It also contained random intercept and slope terms for each individual. A cubic spline fit the data best compared to alternative splines with different degrees-of-freedom and this was assessed using AIC[56]. The internal spline knots were placed at 33.33% and 66.67% quantiles of the data, and boundary knots at the min and max of the data. To assess the significance of diet, the log-likelihood ratio test was used. First, a reduced model excluding the effect of diet was fitted. Then, for both models, the likelihood was computed. Upon computing the ratio of likelihoods, Wilk’s theorem can be used to compute *p*-values to assess significance of the excluded variable (diet in this case). A similar method can be used to compute significance of the diet-time interaction. 95% confidence intervals were estimated from a 1000-sample parametric bootstrap.

Offspring body weight data were analyzed in a similar way as the maternal body weight data. However, the model was changed to include fixed effects of sex and all interactions with both diet as a categorical variable, and a time-on-diet as a cubic natural spline (including three-way interactions). Furthermore, random effects of this model were a quartic natural spline for time-on-diet for each individual. Compared to splines with other degrees-of-freedom, a quartic natural spline was found to be the best model because it had both the minimum AIC[56] and maximum expected-log-predictive-density. The latter was estimated using bayesian hierarchical modelling using the `rstanarm`[57] and `loo`[58] packages. Similar to the maternal body weight data, significance was assessed using the log-likelihood ratio test, and 95% confidence intervals were estimated from a 1000-sample parametric bootstrap. We opted for a maximal model with all interactions so we could make the best possible assessment for the effect of diet[59]. However, we saw similar results with a simpler model: containing fixed effects of growth (modelled as a cubic spline), sex, diet, and only 2-way linear interactions between all variables (i.e. linear growth and sex, linear growth and diet, diet and sex), as well as random effects of growth for each individual (modelled as a cubic spline).

Average body fat percentages in both offspring and dams were analyzed using linear models. An ANOVA was used to assess significance of diet. For maternal data, the linear model had predictors of diet as a categorical variable. For offspring data, the linear model had predictors of diet as a categorical variable, sex, and their interaction. If there was significance in a predictor, a *post hoc* tukey test was done to quantify significance in group differences. Ninety-five percent confidence intervals were also calculated from the model.

Data from glucose tolerance test in both offspring and dams were analyzed using linear mixed-effects models. For maternal data, the model had fixed effects of time (minutes elapsed after IP glucose injection) as a categorical variable with 4 possible values: zero, thirty, sixty, and one-hundred-twenty. These values corresponded to the elapsed time (in minutes) between the IP administration of glucose and when the blood glucose concentration was measured. The model also had fixed effects of diet as a categorical variable and diet-time interactions, as well as a random intercept for each mouse. For the offspring data, the model was similar to the one used with maternal data, but contained an additional fixed effect of sex and all interactions with diet and time (including three-way interactions). Simplifying the model by excluding three-way-interactions did not drastically alter estimates, however, we opted to only present results derived from the maximal model with three-way interactions. Similar to the body weight data, significance of diet was assessed using log-likelihood ratio test. To find specifically which diets were significantly different and at which time points, tukey-adjusted *p*-values were computed using the `emmeans` package[60]. Ninety-five percent confidence intervals were estimated from a 1000-sample parametric bootstrap.

### Neuroanatomy

The log jacobian determinants of the deformation fields from the registration pipeline quantify the volumetric differences between the individual MR image and the consensus average. To assess the effect of diet on neuroanatomy of offspring, a 2-way ANOVA with interaction was conducted on the log determinants at every voxel. The ANOVA had predictors of diet (as a categorical variable), sex, and their interaction. The *F*-statistics associated with diet were extracted for each voxel and converted to *p*-values, which could then be corrected for multiple comparisons using the false discovery rate[61]. Similarly, the *F*-statistics associated with sex-diet interactions were also extracted, converted to *p*-values, and corrected for multiple comparisons.

To visualise the effect of diet on particular structures of interest, labels generated from the MAGeT pipeline were used to find structure volumes. A linear model was fitted to the volume of each structure with predictors of sex, diet (as a categorical variable), and their interaction. This model was then used to define confidence intervals for the various experimental groups. A similar procedure was used to evaluate total brain volume (i.e., sum of all segmented structure volume).

To estimate and visualize the direction of identified effects, we fit a linear-model to the log determinant at each voxel. Each linear model had predictors of sex, dietary fat percentage (as a continuous variable), and their interaction. The *t*-statistics associated with dietary fat percentage effect were extracted from the models, converted to *p*-values, and corrected for multiple comparisons using the false discovery rate[61]. By centering the model on male and female data, *t*-statistics for both sexes were obtained. To visualise the effect of dietary fat percentage on particular structures of interest, the same model was fitted to the structure’s volume.

### Spatial gene expression analysis

Spatial gene expression analysis was conducted similarly to previously published studies using the `ABIgeneRMINC` package[23]. `ANTs`[45] was used to register the Allen Brain Institute (ABI) Atlas space (an average created from serial two-photon images)[22][62] to our MR average. The parameters for registration were a mutual information objective function (32 bins, regular sampling, 0.2 sampling percent) and symmetric normalization transformation (gradient step = 0.1, update field variance = 2, total field variance = 0)[63]. Once the registration was completed, regions-of-interest in MR space were transformed to ABI space for gene expression analysis. To align the developmental gene expression atlas to the MR space, sequential transformations were generated by registering adjacent time points in the ABI gene expression atlas. For example, to align the P14 gene expression atlas to adult brains, the P14 nissl atlas was aligned to the P28 atlas, in turn registered to the adult P56 atlas. The resulting transformations were concatenated to align the P14 gene expression atlas to the P56 atlas, and therefore, the adult MR data.

From the previous voxel-wise analysis, ANOVA was used to quantify the effect of diet on the log determinant at every voxel in the brain. The *F*-statistics associated with diet effects were obtained and transformed to Allen space using the transformation previously described. The top 65% of voxels with the highest *F*-statistics (which corresponds to 1% FDR) constituted the region-of-interest (ROI). We found that our gene expression results were insensitive to this threshold: similar results were seen with an ROI encompassing the top 60% and 70% of voxels. As described by a previous study[23], preferential gene expression was quantified by a fold-change measure: gene expression signal in the ROI divided by gene expression in the whole brain. This fold-change measure was computed for all genes in the ABI gene expression atlas. It is important to note that some genes had incomplete data (i.e. data that did not span a large fraction of the brain). Data from genes where the gene expression signal did not encompass at least 20% of the mouse brain were excluded. Furthermore, if there were multiple datasets encompassing the same gene, only the dataset that provided the largest coverage of the brain was used for the analyses. When clustering the developmental gene expression data, each gene’s preferential expression in the ROI was *Z*−transformed. In order to avoid clustering genes with ubiquitous expression, only genes that had at least 5% increased expression (relative to brain-wide average) at any time point were considered.

Gene Ontology (GO) enrichment analysis was conducted on the top 4000 genes with the greatest preferential expression using the `GOrilla` web application[24], with the background set including all the genes in the mouse genome. To assess significance, `GOrilla` uses the hypergeometric test and corrects for multiple comparisons using false discovery rate[24]. Even though this method of assessing significance is commonly used, the hypergeometric distribution may not be a suitable null distribution as genes and annotations can be correlated[64]. We opted to assess significance using permutation testing. In each of the 10,000 iterations, the association between subjects and diets were shuffled (keeping sex constrained), and the diet-effect on the brain was recalculated. Adult gene expression analysis was re-run on these null brain statistics maps and the top 4000 genes entered into `GOrilla` to obtain a null distribution of GO terms. The fraction of times the FDR values from the null distribution of GO terms was equal to or exceeded the observed FDR value for the GO terms was defined as the *q*-values. GO enrichment analysis yielded similar results when using the top 3000 and 5000 genes, as well as when using a ranked list of genes (i.e. minimum hypergeometric test).

Disease processes associated with gene preferential expression was determined using the Mouse Genome Informatics (MGI) database[65], interfaced using the `InterMineR`[66] package. For all annotated disease terms, the human genes associated with each disease were converted to their homologous mouse genes. Disease terms containing fewer than 15 mouse genes were excluded from the analysis (leaving 13 disease terms in all). Significance of each disease term was assessed using bootstrap. In this procedure, the fold-change of all the disease-associated mouse genes was averaged and this was the test statistic. Then, 10,000 iterations were conducted and, in each iteration, the average fold-change from a random sample of all genes (sampling with replacement) was taken and this defined the null statistic. The *p*-value is defined as the fraction of 10,000 iterations where the null statistic is not less than the test statistic. *p*-values were adjusted for multiple comparisons across the disease terms using bonferroni correction[67]. To validate findings related to autism genes, the Simons Foundation Autism Research Initiative (SFARI) database was used to find genes related to autism mouse models. A bootstrap procedure, identical to the one described previously for MGI database, was used to assess whether these autism-associated genes had significantly high preferential expression. Data from MGI and SFARI was downloaded on April 7, 2020.

## Acknowledgements

This work was supported by grants from the Canadian Institutes of Health Research as well as the Province of Ontario’s Neurodevelopmental Disorders (POND) network of the Ontario Brain Institute. Additional funding support was provided by National Science and Engineering Research Council postgraduate scholarship (DF).

## Supplementary

Exposure to maternal high-fat diet induces extensive changes in the brain of adult offspring. Fernandes *et al*.

**Supplementary Tab. 1a:** LF10 Rodent Diet (Research Diets D12450Bi) Formulation.

| Class description | Ingredient | Grams (g) |
| --- | --- | --- |
| Protein | Casein, Lactic, 30 Mesh | 200.00 |
| Protein | Cystine, L | 3.00 |
| Carbohydrate | Sucrose, fine granulated | 354.00 |
| Carbohydrate | Starch, corn | 315.00 |
| Carbohydrate | Lodex 10 | 35.00 |
| Fiber | Solka Floc, FCC200 | 50.00 |
| Fat | Lard | 20.00 |
| Fat | Soybean oil, USP | 25.00 |
| Mineral | S10026B | 50.00 |
| Vitamin | Choline bitartrate | 2.00 |
| Vitamin | V10001C | 1.00 |
| Dye | Dye, Yellow FD&C no.5, Alum. Lake 35-42% | 0.05 |
|  | **Total** | 1055.05 |

**Supplementary Tab. 1b:** HF45 Rodent Diet (Research Diets D12451i) Formulation.

| Class description | Ingredient | Grams (g) |
| --- | --- | --- |
| Protein | Casein, Lactic, 30 Mesh | 200.00 |
| Protein | Cystine, L | 3.00 |
| Carbohydrate | Sucrose, fine granulated | 176.80 |
| Carbohydrate | Starch, corn | 72.80 |
| Carbohydrate | Lodex 10 | 100 |
| Fiber | Solka Floc, FCC200 | 50.00 |
| Fat | Lard | 177.50 |
| Fat | Soybean oil, USP | 25.00 |
| Mineral | S10026B | 50.00 |
| Vitamin | Choline bitartrate | 2.00 |
| Vitamin | V10001C | 1.00 |
| Dye | Dye, Red FD&C no.40, Alum. Lake 35-42% | 0.05 |
|  | **Total** | 858.15 |

**Supplementary Tab. 1c:** HF60 Rodent Diet (Research Diets D12492i) Formulation.

| Class description | Ingredient | Grams (g) |
| --- | --- | --- |
| Protein | Casein, Lactic, 30 Mesh | 200.00 |
| Protein | Cystine, L | 3.00 |
| Carbohydrate | Sucrose, fine granulated | 72.80 |
| Carbohydrate | Lodex 10 | 125.00 |
| Fiber | Solka Floc, FCC200 | 50.00 |
| Fat | Lard | 245.00 |
| Fat | Soybean oil, USP | 25.00 |
| Mineral | S10026B | 50.00 |
| Vitamin | Choline bitartrate | 2.00 |
| Vitamin | V10001C | 1.00 |
| Dye | Dye, Blue FD&C no.1, Alum. Lake 35-42% | 0.05 |
|  | **Total** | 773.85 |

**Supplementary Tab. 2:** Diet Caloric Information and Fuel Values.

| Diet | Protein (% kcal) | Fat (% kcal) | Carbohydrate (% kcal) | Energy Density (kcal/g) |
| --- | --- | --- | --- | --- |
| LF10 | 20 | 10 | 70 | 3.82 |
| HF45 | 20 | 45 | 35 | 4.7 |
| HF60 | 20 | 60 | 20 | 5.21 |

## References

1. Stubert, J., Reister, F., Hartmann, S. & Janni, W. The risks associated with obesity in pregnancy. Deutsches Ärzteblatt International 115, 276 (2018).

2. Cedergren, M. I. Maternal morbid obesity and the risk of adverse pregnancy outcome. Obstetrics & Gynecology 103, 219–224 (2004).

3. Parlee, S. D. & MacDougald, O. A. Maternal nutrition and risk of obesity in offspring: the Trojan horse of developmental plasticity. Biochimica et Biophysica Acta (BBA)-Molecular Basis of Disease 1842, 495–506 (2014).

4. Reynolds, R. M. et al. Maternal obesity during pregnancy and premature mortality from cardiovascular event in adult offspring: follow-up of 1 323 275 person years. Bmj 347, f4539 (2013).

5. Grattan, D. R. Fetal programming from maternal obesity: eating too much for two? Endocrinology 149, 5345–5347 (2008).

6. Bilder, D. A. et al. Maternal prenatal weight gain and autism spectrum disorders. Pediatrics 132, e1276–e1283 (2013).

7. Toth, M. Mechanisms of non-genetic inheritance and psychiatric disorders. Neuropsychopharmacology 40, 129–140 (2015).

8. Brown, A. S. Epidemiologic studies of exposure to prenatal infection and risk of schizophrenia and autism. Developmental neurobiology 72, 1272–1276 (2012).

9. Mina, T. H. et al. Prenatal exposure to very severe maternal obesity is associated with adverse neuropsychiatric outcomes in children. Psychological medicine 47, 353–362 (2017).

10. Robinson, M. et al. Pre-pregnancy maternal overweight and obesity increase the risk for affective disorders in offspring. Journal of developmental origins of health and disease 4, 42–48 (2013).

11. Rivera, H. M., Christiansen, K. J. & Sullivan, E. L. The role of maternal obesity in the risk of neuropsychiatric disorders. Frontiers in neuroscience 9, 194 (2015).

12. Sullivan, E. L., Riper, K. M., Lockard, R. & Valleau, J. C. Maternal high-fat diet programming of the neuroendocrine system and behavior. Hormones and behavior 76, 153–161 (2015).

13. Kang, S. S., Kurti, A., Fair, D. A. & Fryer, J. D. Dietary intervention rescues maternal obesity induced behavior deficits and neuroinflammation in offspring. Journal of neuroinflammation 11, 156 (2014).

14. Kleinert, M. et al. Animal models of obesity and diabetes mellitus. Nature Reviews Endocrinology 14, 140 (2018).

15. Wang, C.-Y. & Liao, J. K. in mTOR 421–433 (Springer, 2012).

16. Can, Ö. D., Ulupinar, E., Özkay, Ü. D., Yegin, B. & Öztürk, Y. The effect of simvastatin treatment on behavioral parameters, cognitive performance, and hippocampal morphology in rats fed a standard or a high-fat diet. Behavioural pharmacology 23, 582–592 (2012).

17. Bilbo, S. D. & Tsang, V. Enduring consequences of maternal obesity for brain inflammation and behavior of offspring. The FASEB Journal 24, 2104–2115 (2010).

18. Giriko, C. Á. et al. Delayed physical and neurobehavioral development and increased aggressive and depression-like behaviors in the rat offspring of dams fed a high-fat diet. International journal of developmental neuroscience 31, 731–739 (2013).

19. Sasaki, A., De Vega, W., St-Cyr, S., Pan, P. & McGowan, P. Perinatal high fat diet alters glucocorticoid signaling and anxiety behavior in adulthood. Neuroscience 240, 1–12 (2013).

20. Tozuka, Y. et al. Maternal obesity impairs hippocampal BDNF production and spatial learning performance in young mouse offspring. Neurochemistry international 57, 235–247 (2010).

21. Dazai, J., Spring, S., Cahill, L. S. & Henkelman, R. M. Multiple-mouse neuroanatomical magnetic resonance imaging. JoVE (Journal of Visualized Experiments), e2497 (2011).

22. Lein, E. S. et al. Genome-wide atlas of gene expression in the adult mouse brain. Nature 445, 168 (2007).

23. Fernandes, D. J. et al. Spatial gene expression analysis of neuroanatomical differences in mouse models. Neuroimage 163, 220–230 (2017).

24. Eden, E., Navon, R., Steinfeld, I., Lipson, D. & Yakhini, Z. GOrilla: a tool for discovery and visualization of enriched GO terms in ranked gene lists. BMC bioinformatics 10, 48 (2009).

25. Yang, Y. et al. Early-life high-fat diet-induced obesity programs hippocampal development and cognitive functions via regulation of gut commensal Akkermansia muciniphila. Neuropsychopharmacology 44, 2054–2064 (2019).

26. Teegarden, S. L., Scott, A. N. & Bale, T. L. Early life exposure to a high fat diet promotes long-term changes in dietary pREFerences and central reward signaling. Neuroscience 162, 924–932 (2009).

27. Hariri, N. & Thibault, L. High-fat diet-induced obesity in animal models. Nutrition research reviews 23, 270–299 (2010).

28. Huang, X.-F. et al. Effects of diets high in whey, soy, red meat and milk protein on body weight maintenance in diet-induced obesity in mice. Nutrition & dietetics 65, S53–S59 (2008).

29. Ellegood, J. et al. Clustering autism: using neuroanatomical differences in 26 mouse models to gain insight into the heterogeneity. Molecular psychiatry 20, 118–125 (2015).

30. Rollins, C. P. et al. Contributions of a high-fat diet to Alzheimer’s disease-related decline: A longitudinal behavioural and structural neuroimaging study in mouse models. NeuroImage: Clinical 21, 101606 (2019).

31. Sims-Robinson, C. et al. Dietary reversal ameliorates short-and long-term memory deficits induced by high-fat diet early in life. PloS one 11 (2016).

32. Koob, G. F. Neurobiology of addiction: toward the development of new therapies. Annals of the New York Academy of Sciences 909, 170–185 (2000).

33. Kenny, P. J. Reward mechanisms in obesity: new insights and future directions. Neuron 69, 664–679 (2011).

34. Unal, C. T., Golowasch, J. P. & Zaborszky, L. Adult mouse basal forebrain harbors two distinct cholinergic populations defined by their electrophysiology. Frontiers in behavioral neuroscience 6, 21 (2012).

35. De Jonge, F., Oldenburger, W., Louwerse, A. & Van de Poll, N. E. Changes in male copulatory behavior after sexual exciting stimuli: effects of medial amygdala lesions. Physiology & behavior 52, 327–332 (1992).

36. Spring, S., Lerch, J. P. & Henkelman, R. M. Sexual dimorphism revealed in the structure of the mouse brain using three-dimensional magnetic resonance imaging. Neuroimage 35, 1424–1433 (2007).

37. Sáez-López, C., Rivera-Giméenez, M., Hernéandez, C., Siméo, R. & Selva, D. M. SHBG-C57BL/ksJ-db/db: a new mouse model to study SHBG expression and regulation during obesity development. Endocrinology 156, 4571–4581 (2015).

38. Curtis, D. & Consortium, U. Practical experience of the application of a weighted burden test to whole exome sequence data for obesity and schizophrenia. Annals of human genetics 80, 38–49 (2016).

39. Corre, C. et al. Sex-specific regulation of weight and puberty by the Lin28/let-7 axis. The Journal of endocrinology 228, 179 (2016).

40. Grasemann, C. et al. Parental diabetes: the Akita mouse as a model of the effects of maternal and paternal hyperglycemia in wildtype offspring. PLoS One 7 (2012).

41. Cahill, L. S. et al. Preparation of fixed mouse brains for MRI. Neuroimage 60, 933–939 (2012).

42. Spencer Noakes, T. L., Henkelman, R. M. & Nieman, B. J. Partitioning k-space for cylindrical three-dimensional rapid acquisition with relaxation enhancement imaging in the mouse brain. NMR in biomedicine 30, e3802 (2017).

43. Lerch, J. P. et al. Maze training in mice induces MRI-detectable brain shape changes specific to the type of learning. Neuroimage 54, 2086–2095 (2011).

44. Collins, D. L., Neelin, P., Peters, T. M. & Evans, A. C. Automatic 3D intersubject registration of MR volumetric data in standardized Talairach space. Journal of computer assisted tomography 18, 192–205 (1994).

45. Avants, B. B., Tustison, N. & Song, G. Advanced normalization tools (ANTS). Insight j 2, 1–35 (2009).

46. Friedel, M., van Eede, M. C., Pipitone, J., Chakravarty, M. M. & Lerch, J. P. Pydpiper: a flexible toolkit for constructing novel registration pipelines. Frontiers in neuroinformatics 8, 67 (2014).

47. Chakravarty, M. M. et al. Performing label-fusion-based segmentation using multiple automatically generated templates. Human brain mapping 34, 2635–2654 (2013).

48. Dorr, A., Lerch, J. P., Spring, S., Kabani, N. & Henkelman, R. M. High resolution three-dimensional brain atlas using an average magnetic resonance image of 40 adult C57Bl/6J mice. Neuroimage 42, 60–69 (2008).

49. Steadman, P. E. et al. Genetic effects on cerebellar structure across mouse models of autism using a magnetic resonance imaging atlas. Autism research 7, 124–137 (2014).

50. Ullmann, J. F., Watson, C., Janke, A. L., Kurniawan, N. D. & Reutens, D. C. A segmentation protocol and MRI atlas of the C57BL/6J mouse neocortex. Neuroimage 78, 196–203 (2013).

51. Richards, K. et al. Segmentation of the mouse hippocampal formation in magnetic resonance images. Neuroimage 58, 732–740 (2011).

52. Qiu, L. R. et al. Mouse MRI shows brain areas relatively larger in males emerge before those larger in females. Nature communications 9, 2615 (2018).

53. Wickham, H. tidyverse: Easily Install and Load the ‘Tidyverse’ R package version 1.2.1 (2017). https://CRAN.R-project.org/package=tidyverse.

54. Kuznetsova, A., Brockhoff, P. B. & Christensen, R. H. B. lmerTest Package: Tests in Linear Mixed Effects Models. Journal of Statistical Software 82, 1–26 (2017).

55. Bates, D., Mächler, M., Bolker, B. & Walker, S. Fitting Linear Mixed-Effects Models Using lme4. Journal of Statistical Software 67, 1–48 (2015).

56. Burnham, K. P. & Anderson, D. R. A practical information-theoretic approach. Model selection and multimodel inference, 2nd ed. Springer, New York (2002).

57. Stan Development Team. rstanarm: Bayesian applied regression modeling via Stan. R package version 2.13.1. 2016. http://mc-stan.org/.

58. Vehtari, A., Gabry, J., Yao, Y. & Gelman, A. loo: Efficient leave-one-out cross-validation and WAIC for Bayesian models R package version 2.0.0. 2018. https://CRAN.R-project.org/package=loo.

59. Barr, D. J., Levy, R., Scheepers, C. & Tily, H. J. Random effects structure for confirmatory hypothesis testing: Keep it maximal. Journal of memory and language 68, 255–278 (2013).

60. Lenth, R. emmeans: Estimated Marginal Means, aka Least-Squares Means R package version 3.5.1 (2019). https://CRAN.R-project.org/package=emmeans.

61. Genovese, C. R., Lazar, N. A. & Nichols, T. Thresholding of statistical maps in functional neuroimaging using the false discovery rate. Neuroimage 15, 870–878 (2002).

62. Oh, S. W. et al. A mesoscale connectome of the mouse brain. Nature 508, 207 (2014).

63. Yee, Y. et al. Structural covariance of brain region volumes is associated with both structural connectivity and transcriptomic similarity. Neuroimage 179, 357–372 (2018).

64. Goeman, J. J. & Bühlmann, P. Analyzing gene expression data in terms of gene sets: methodological issues. Bioinformatics 23, 980–987 (2007).

65. Motenko, H., Neuhauser, S. B., O’keefe, M. & Richardson, J. E. MouseMine: a new data warehouse for MGI. Mammalian Genome 26, 325–330 (2015).

66. Kyritsis, K. A., Wang, B., Sullivan, J., Lyne, R. & Micklem, G. InterMineR: an R package for InterMine databases. Bioinformatics 35, 3206–3207 (2019).

67. Dunn, O. J. Multiple comparisons among means. Journal of the American statistical association 56, 52–64 (1961).

